# Redox homeostasis in ferroptosis and aging: a causal role for *fard-1* and *dhs-25* in *Caenorhabditis elegans*

**DOI:** 10.1101/2025.01.07.631721

**Authors:** Roberta Pensotti, Barbara Sciandrone, Federica Bovio, Laura Schröter, Silvia Maglioni, Leonie Thewes, Matilde Emma Forcella, Paola Alessandra Fusi, Andrea Rossi, Carsten Berndt, Natascia Ventura, Maria Elena Regonesi

**Author notes:** These authors co-last authored this manuscript (Natascia Ventura, Maria Elena Regonesi).

## Abstract

Aging is a natural process characterized by a progressive physiological decline that undermines health and well-being in the elderly population. It is widely accepted that an unbalanced redox state belongs to the hallmarks of aging, but its role as one of the main drivers of ferroptosis is quite recent. Ferroptosis is a form of iron-dependent cell death caused by massive phospholipid peroxidation. The excessive accumulation of intracellular reactive oxygen species and iron, as well as the failure of the main cellular antioxidant systems, cause ferroptotic cell death. While clear roles for ferroptosis in pathological conditions such as cancer or neurodegeneration have been described, its physiological roles and regulators are less clearly understood.

Here, using *Caenorhabditis elegans* as a powerful model organism for aging studies, we uncover a role for ferroptosis in physiological aging mediated by disturbed redox homeostasis. We evaluated healthspan parameters in a *C. elegans* wild-type strain highlighting how several age-related features differentially decline during aging. A progressive loss of the capability to contrast external stressors, with an increase in hydroxyl radicals and a failure of the glutathione antioxidant system demonstrated the progressive disruption of redox homeostasis in older age. Moreover, we showed that selected genes involved in redox metabolism are downregulated with aging. Among them, mutant strains of the fatty acyl-CoA reductase, *fard-1*, and of the dehydrogenase, *dhs-25*, displayed higher sensitivity to a ferroptosis inducer, increased lipid peroxidation, anticipated drop in total glutathione and reduced lifespan. Accordingly, the expression of one of the closest mammalians *dhs-25* homolog, the hydroxysteroid 17-Beta Dehydrogenase 8, was downregulated in cells which are more sensitive to ferroptosis.

Our results clearly prove a causal role for ferroptosis in *C. elegans* aging driven by oxidative stress, unveiling novel genes involved in this connection that may constitute targets for possible interventions to improve healthy aging.

## INTRODUCTION

One of the greatest challenges of the 21st century is the progressively aging society: people over 60 years old are expected to reach more than 20% of the global population by 2050. This increase in life expectancy is related to the modern medical and technological improvements that have led to the identification of novel therapeutic approaches against fatal disorders, thus resulting in a reduced mortality rate in many nations around the world ^1^. Despite these positive achievements, the current transition towards an aging population implies several disadvantages for the society and the elderly themselves. Indeed, the aging process is characterized by a progressive time-dependent decline in physiological functions that leads to increased frailty and a growing risk of age-related disorders (e.g. cancer, diabetes, cardiovascular and neurodegenerative disorders) ^2–5^. The interplay between twelve hallmarks characterizes this process: genomic instability, telomere attrition, epigenetic alteration, loss of proteostasis, altered macroautophagy, deregulated nutrient-sensing, mitochondrial dysfunction, cellular senescence, stem cell exhaustion, altered intracellular communication, chronic inflammation and dysbiosis ^6,7^. In particular, mitochondria undergo a progressive deterioration with age showing an increased production of reactive oxygen species (ROS), thus leading to an unbalanced redox homeostasis that spreads throughout the cell ^8–11^. Oxidative stress has recently been identified as one of the main drivers of ferroptosis, a newly discovered iron-dependent form of cell death caused by the PUFA-phospholipids peroxidation, which represents its distinctive hallmark ^12–14^. In detail, the progressive accumulation of phospholipid peroxidation increases the membrane tension and challenges its ionic and osmotic homeostasis, up to the inexorable rupture of the lipid bilayer and the consequent cell death ^15–17^. Other important factors that trigger the ferroptotic process are the high levels of free intracellular iron in the Fe^2+^ form, that promote the initiation and propagation of lipid peroxidation through the Fenton reaction, and the failure of the antioxidant systems responsible for the PUFA-peroxides detoxification ^18,19^. Among these, the main antioxidant systems involved in contrasting ferroptosis include glutathione peroxidase 4 (GPX4) and ferroptosis suppressor protein 1 (FSP1). The GPX4 plays a key role by converting the toxic phospholipid hydroperoxides (PLOOHs) into non-toxic phospholipid alcohols (PLOHs), using two electrons provided by the reduced form of glutathione (GSH) ^20–22^. Instead, FSP1 is a NAD(P)H-dependent oxidoreductase, localized in the cell membrane, responsible for coenzyme Q10 (CoQ10) and vitamin K reduction ^23–25^. These species, acting as lipophilic radical-trapping antioxidants, can directly reduce lipid radicals, thus stopping the spread of lipid peroxidation ^26^. Therefore, multiple cellular metabolic pathways are associated with ferroptosis, highlighting its implication in the progression of different pathologies, including infectious diseases, neurodegeneration and cancer ^27–31^. In particular, it has been observed that a variety of cancer cells resistant to traditional therapies or with a high propensity to metastasize are more susceptible to ferroptosis-inducing drugs ^32–34^. Moreover, a recent study reported that human α-synuclein aggregates activate ferroptosis, contributing to parvalbumin interneuron degeneration and motor circuit dysfunction ^35^. Despite its implication in tumour suppression and immune surveillance ^36^, its physiological relevance remains unclear. Recently, some studies carried out in *Caenorhabditis elegans* model have proven that ferroptosis is an evolutionarily conserved process that contributes to accelerate the aging process. Schiavi et al. showed that silencing frataxin, a nuclear-encoded mitochondrial protein involved in ISC (Fe-S clusters)’ biogenesis, leads to lifespan extension in *C. elegans* through the inhibition of ferroptosis as one of the possible mechanisms ^37^. Moreover, Jenkins et al. have recently demonstrated that the combination of increased levels of labile iron and depletion of glutathione, during normal aging, correlate with increased risk of ferroptosis, enhanced organism frailty and shortened *C. elegans* lifespan ^38^. The use of *C. elegans* as a model organism in the aging field is widely established. The short lifespan (maximum 3 weeks at 20 °C), the presence of >40% of coding genes that are mammalian orthologs ^39^, the easy maintenance and manipulation that allow reproducible and high-throughput experiments, and the availability of molecular biology tools (i.e., transgenic strains ^40^, gene knockouts ^41^, and RNAi knockdowns ^42^) make this nematode a unique model to study aging. Moreover, many mechanisms regulating longevity, including nutrient sensing, redox balance and protein homeostasis, are evolutionally conserved from worms to mammals ^43,44^. Noteworthy, exploiting these advantages, several breakthrough discoveries in aging research have been achieved using *C. elegans*, which have been subsequently confirmed in higher organisms.

In this work we employed *C. elegans* to investigate the implication of ferroptosis in the aging process but also to identify new possible ferroptosis regulatory genes exploitable as targets for limiting frailty and improving healthy aging. At first, extensive phenotypical characterization of different healthspan parameters (movement, pumping rate, oxidative and thermal stress resistance) during *C. elegans* aging allowed to identify three key time points (day 4, 7 and 14 of adult life). At these time points, biochemical and molecular parameters of relevance for redox homeostasis were quantified, namely the formation of hydroxyl radicals, the total glutathione amount with the ratio of its reduced/oxidized forms, and the expression of redox-associated genes. Oxidative stress condition, lifespan, lipid peroxidation and sensitivity to ferroptosis were then analysed in strains knocked out for those genes which resulted to be downregulated during aging. Our analysis identified two genes with oxidoreductase activity, *fard-1* and *dhs-25*, causally involved in the ferroptosis-aging association, with evolutionarily conserved implication for the *dhs-25* mammalian ortholog in ferroptosis.

## RESULTS

### Identification of critical health time points during *C. elegans* lifespan

Aging is an evolutionarily conserved process characterized by progressive decline of different physiological features ^6^. Accordingly, during *C. elegans* aging there is a progressive decline of the main healthspan parameters i.e. locomotion, pumping rate, resistance to stress, which nonetheless may change at different peace ^45,46^. To identify critical life points, different *C. elegans* health-related parameters were closely monitored during lifespan under standard experimental conditions, using wild-type strain maintained on nematode growth media (NGM) agar plates at 20 °C seeded with alive *E. coli* OP50 at the final concentration of OD_600_=0.3 as a food source. Kaplan-Meier survival analyses (Figure 1a) revealed a median (day 14) and maximum (day 19) lifespan value consistent with the literature ^47^. Animals’ motility was evaluated during lifespan by counting body bends in a large population of ∼1 000 worms starting from the first day of adulthood as day 0. Despite high variability in the counts within the selected cohort, as highlighted by the box plot representation, a general linear decline in locomotion was observed starting from early adulthood (Figure 1b). Feeding rate was measured by pharyngeal pumping assay in a total number of 90 worms per selected time point of lifespan (Figure 1c). Similar to locomotion, pharyngeal contractions decreased in a progressive linear manner starting from young adult animals. Lastly, the ability of the nematodes to counteract external stressors (i.e. thermal and oxidative stresses) was assessed. Resistance to thermal stress represents one of the most reliable surrogate marker and predictor of lifespan ^48^. Thermotolerance test was carried out in *C. elegans* by laying the animals at high temperatures. Nematodes live at ∼15-20 °C as optimum, whereas exposure to temperatures above 25 °C and especially between 30 and 37 °C induces thermal stress, leading to the progressive accumulation of cellular damages and a shortened lifespan (survival tests can be completed within 24 hours) ^49–51^. Starting from a synchronous population at day 0, 30 animals were scored for survival at 37 °C every two days until day 16 of adulthood (Figure 1d). The results clearly showed that the ability to withstand heat shock did not change in the first week of life but began to decline from the second week (Figure 1d). Similarly, the median survival of 30 worms exposed to hydrogen peroxide, used as oxidative stressor, remained unchanged in the first week (Figure 1e). Unfortunately, reliable data could not be obtained from the second week of lifespan possibly due to the significant reduction in hydrogen peroxide uptake, caused by cuticle thickening during aging ^52^, as well as the substantial decline in pharyngeal pumping rate observed above (Figure 1c).

**Figure 1.**
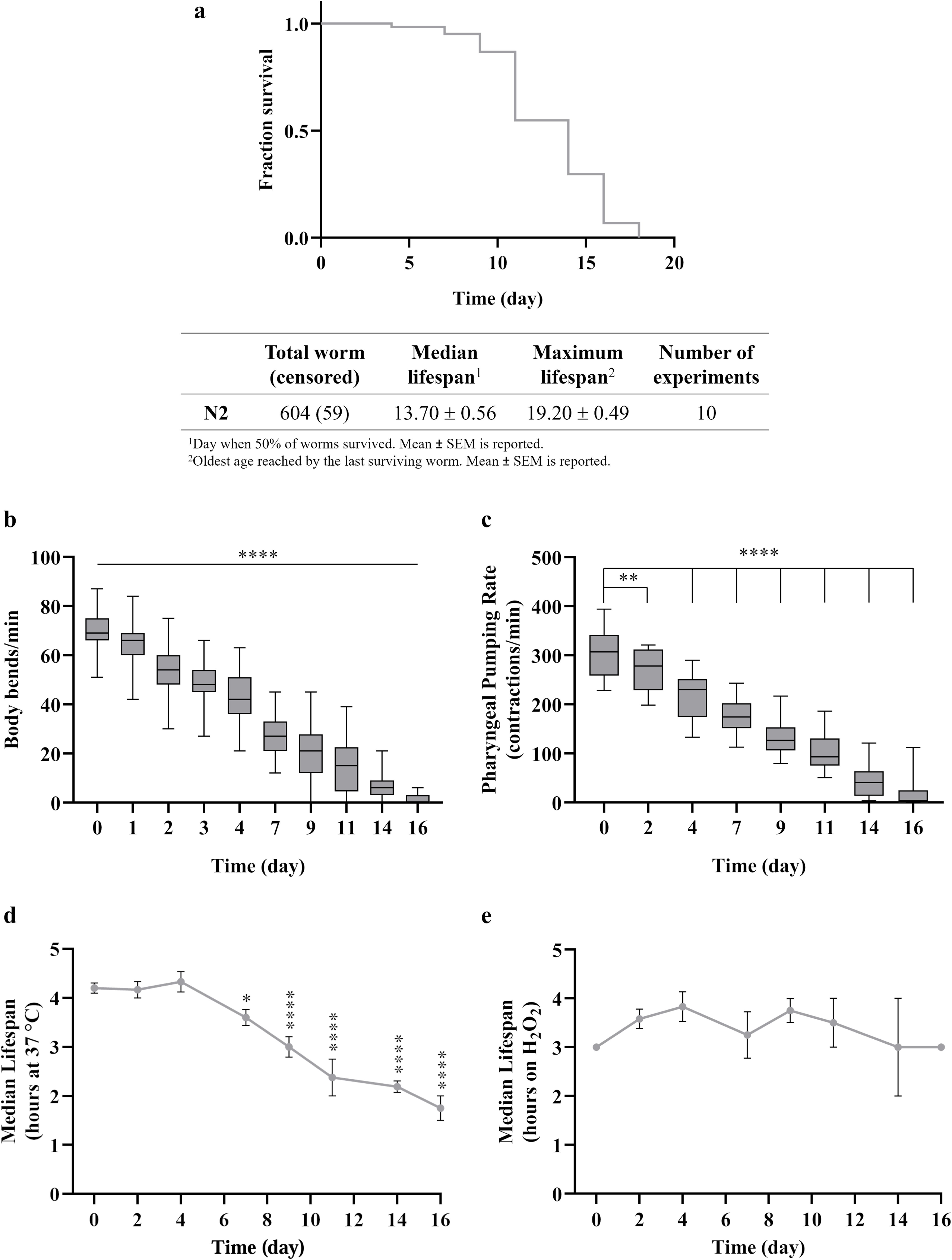
Characterization of healthspan changes during lifespan in a wild-type N2 *C. elegans* strain. (a). Survival analysis were performed on a population of synchronous 1-day adult worms placed on NGM plates at 20 °C with alive *E. coli* OP50 OD_600_=0.3 as food cource. A representative Kaplan-Meier curve carried out with 60 animals is shown and the median and maximum days of lifespan are expressed as mean ± SEM from ten independent experiments. (b) Locomotion of a wide population of worms (∼ 1 000) was assessed over time. The starting population was mantained by tranferring the worms on fresh plates every day during the fertile period and every two days afterwards. The body bends of each single worm were counted at days 0, 1, 2, 3, 4, 7, 9, 11, 14, 16. Collected data were assembled in a box and whisker plot to display how values are dispersed. (c) The pharyngeal pumps of 90 worms were counted at days 0, 2, 4, 7, 9, 11, 14, 16 and assembled in box and whiskers. (d) Survival at 37 °C of 30 worms was scored at days 0, 2, 4, 7, 9, 11, 14, 16. Median survival at 37 °C for each time point of life was plotted as mean ± SEM of at least three independent experiments. (e) Survival of 30 animals in the presence of 0.03% H_2_O_2_ was scored at days 0, 2, 4, 7, 9, 11, 14, 16. Median survival to H_2_O_2_ was plotted as mean ± SEM of three independent experiments. *p ≤ 0.05; **p ≤ 0.01; ****p ≤ 0.0001 by one-way ANOVA with Dunnett’s multiple comparisons test.

Overall, three main points of change (day 4, 7, and 14) were identified and selected for subsequent experiments, considering day 0, the1^st^ day of adulthood, as a reference.

### Failure of redox homeostasis during *C. elegans* lifespan

One of the main hallmarks of aging is the failure of redox homeostasis, characterized by increased production of oxidants and accumulation of free iron, coupled to the decline of the antioxidant defenses ^53–55^. Therefore, the amount of hydroxyl radicals produced by the Fenton reaction was measured in worm’ lysates collected at day 0, 4, 7 and 14 of adulthood by fluorometric assay using 2’,7’-dichlorodihydrofluorescein-diacetate probe. Collected data clearly showed a time-dependent increase in Fenton reaction which became statistically significant starting from day 7 and peaked at day 14 (Figure 2a). Since glutathione is one of the main antioxidant systems conserved from worms to mammals ^56^, its total levels and the ratio of reduced to oxidized forms (GSH/GSSG) were determined using a colorimetric assay based on 5,5′-dithiobis(2-nitrobenzoic acid) reduction, at the same time points. Contrary to our expectation, total glutathione levels exhibited a progressive increase that became significant by day 7 (Figure 2b), possibly reflecting a compensatory response to rising oxidative damage. This increase was nonetheless followed by a dramatic drop at day 14 (Figure 2b), indicating the inability of the animals to compensate the higher oxidative stress at this later stage of life. Consistent with this, the GSH/GSSG ratio switched progressively from 0.69 at day 0 to 2.05 by day 14 (Figure 2c), as expected by the progressive increase in oxidative environment during aging.

**Figure 2.**
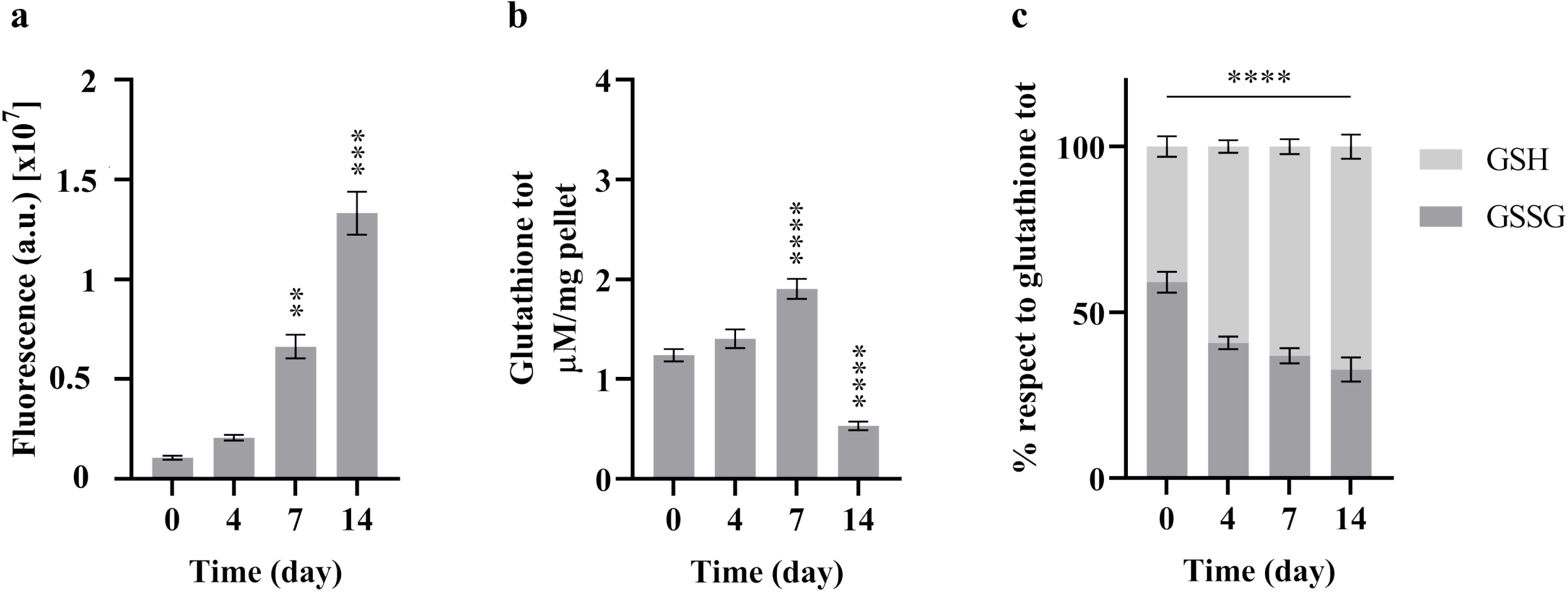
Characterization of changes in redox homeostasis during wild-type N2 *C. elegans* aging. (a) Hydroxyl radicals were quantified measuring the fluorescence intensity of the oxidized form of H_2_DCFDA in lysates of nematodes at day 0, 4, 7, 14 of adulthood. A starting population of worms was maintained by transferring the animals on fresh plates every day during the fertile period and every two days afterwards. At each specified time point, 100 worms were collected. Fluorescence intensity in arbitrary units was reported as mean ± SEM of at least three replicates. (b) Total amount of glutathione was derived by DTNB reduction colorimetric assay in lysates of nematodes at day 0, 4, 7, 14 of adulthood. At each specified time point, around 400 worms were collected from a starting population maintained over time. Normalized values of glutathione concentration (μM/mg) were reported as mean ± SEM of at least three biological replicates. (c) The incubation of 100 μL of worms’ lysate, processed as in b, with the GSH masking reagent 2-vinylpyridine allowed to determine the GSH/GSSG ratio. The percentage values of GSH and GSSG respect to total glutathione for each time point were represented as mean percentage distribution ± % SEM of at least three biological replicates. **p ≤ 0.01; ***p ≤ 0.001; ****p ≤ 0.0001 by one-way ANOVA with Dunnett’s multiple comparisons test.

### The expression of redox-associated genes is significantly decreased during *C. elegans* aging

In search of genes which might help counteracting the oxidative environment during aging, we turned to differentially expressed genes (DEG) previously identified in long-lived animals with mild mitochondrial stress due to partial frataxin (*frh-1*) depletion ^37^. Silencing of frataxin, a mitochondrial protein involved in iron homeostasis and ISC biogenesis, was indeed demonstrated to extend worms’ lifespan through inhibition of ferroptosis ^37^. This prompt us to speculate that DEGs in long-lived *frh-1*-depleted worms could be involved in the regulation of physiological aging via redox-regulated ferroptosis. Genes with oxidoreductase activity induced by *frh-1* RNAi were thus selected (Table 1; Table S1) and their expression was first validated in *frh-1* silenced animals (Figure S1) and then investigated during wild-type, physiological aging. The results displayed that most of the genes that were upregulated in *frh-1* RNAi worms were subjected to downregulation during lifespan in a wild-type background (Figure 3a-b, i-k, m-s, u), confirming their possible implication in the aging and ferroptotic processes. Some exceptions included *trx-2* (Figure 3t) and *dhs-18* (Figure 3h), whose mRNA levels slightly increased at day 4 and 7; *ctl-3* (Figure 3c), *cyp-13A8* (Figure 3d) and *cyp-33C8* (Figure 3g) genes that were upregulated during aging; and *cyp-14A1* (Figure 3e) and *cyp-14A4* (Figure 3f) which were unchanged over time. The only selected downregulated gene in *frh-1* RNAi worms, *fat-6*, after an initial upregulation at day 4 and 7 of adulthood, returned to expression levels comparable to young animals (Figure 3l). Of note, human orthologs of some of the genes upregulated by *frh-1* RNAi and downregulated during physiological aging, have been already involved in the ferroptosis process, such the GPX ortholog *gpx-7* (Figure 3o).

**Figure 3.**
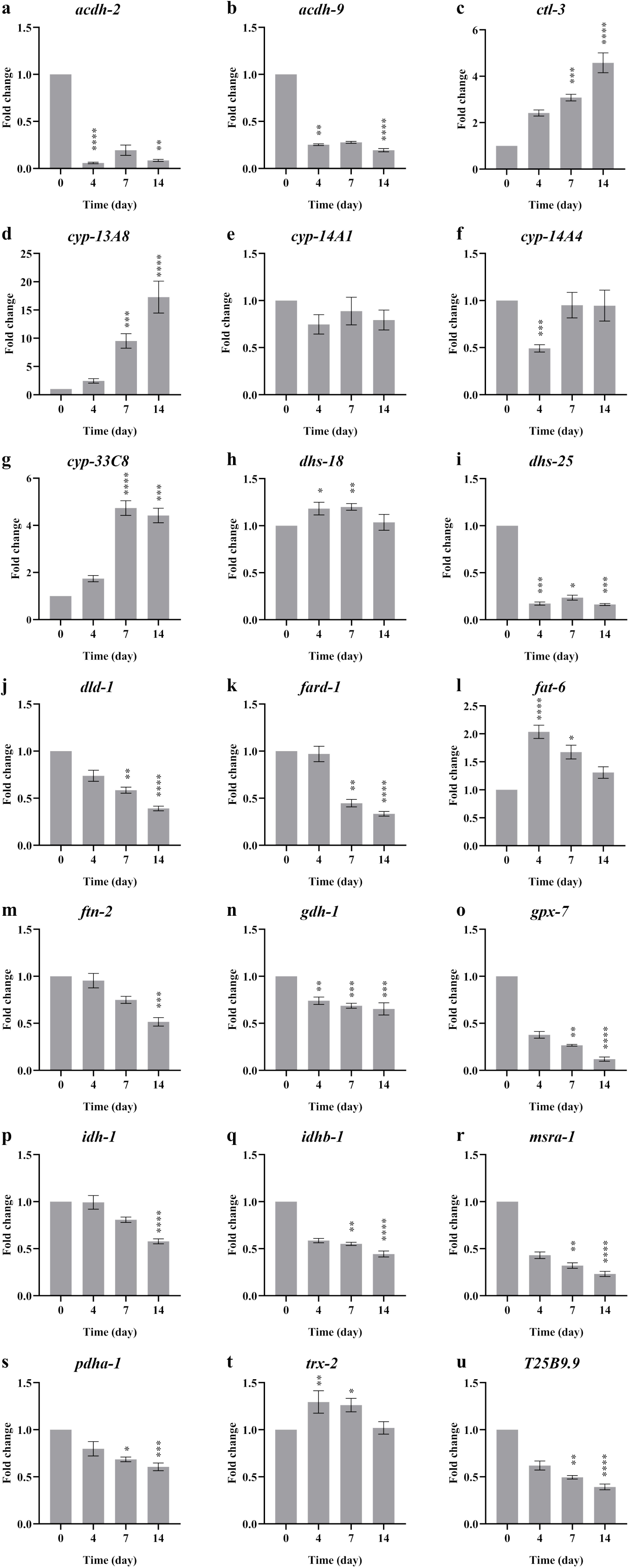
Evaluation of the selected redox-genes expression during N2 aging. Real Time PCR analysis of the genes selected from *frh-1* RNAi worms trascriptomic analysis were carried out on worms collected at day 0, 4, 7, 14 of adulthood. Around 1 000-2 000 nematodes for each time point were collected for RNA extraction and reverse transcription. *cdc-42* was chosen as housekeeping gene for the normalization. The fold change value of each gene was reported as mean ± SEM of the biological triplicate. (a) *acdh-2*, (b) *acdh-9*, (c) *ctl-3*, (d) *cyp-13A8*, (e) *cyp-14A1*, (f) *cyp-14A4*, (g) *cyp-33C8*, (h) *dhs-18*, (i) *dhs-25*, (j) *dld-1*, (k) *fard-1*, (l) *fat-6*, (m) *ftn-2*, (n) *gdh-1*, (o) *gpx-7*, (p) *idh-1*, (q) *idhb-1*, (r) *msra-1*, (s) *pdha-1*, (t) *trx-2*, (u) *T25B9.9*. *p ≤ 0.05; **p ≤ 0.01; ***p ≤ 0.001; ****p ≤ 0.0001 by Kruskal-Wallis, Dunn’s multiple comparisons test in GraphPad Prism 8.0.2.

**Table 1.** Description of DEG genes selected in *frh-1* RNAi worms with related fold changes.

| Gene | Gene name | Fold change in *frh-1* RNAi worms |
| --- | --- | --- |
| *acdh-2* | Acyl CoA dehydrogenase | 1.420 |
| *acdh-9* | Acyl CoA dehydrogenase | 0.956 |
| *cdc-42* | Cell division cycle related | N.A. |
| *ctl-3* | Catalase | 0.436 |
| *cyp-13A8* | Cytochrome P450 family | 5.304 |
| *cyp-14A1* | Cytochrome P450 family | 2.916 |
| *cyp-14A4* | Cytochrome P450 family | 4.907 |
| *cyp-33C8* | Cytochrome P450 family | 3.934 |
| *dhs-18* | Dehydrogenases, short chain | 0.479 |
| *dhs-25* | Dehydrogenases, short chain | 2.372 |
| *dld-1* | Dihydrolipoamide dehydrogenase | 0.314 |
| *fard-1* | Fatty acyl-CoA reductase | 0.991 |
| *fat-6* | Fatty acid desaturase | -0.403 |
| *ftn-2* | Ferritin | -0.479 |
| *gdh-1* | Glutamate dehydrogenase | 0.269 |
| *gpx-7* | Glutathione peroxidase | 0.442 |
| *idh-1* | Isocitrate dehydrogenase | 0.395 |
| *idhb-1* | Isocitrate dehydrogenase beta | 0.277 |
| *msra-1* | Methionine sulfoxide reductase A | 0.696 |
| *pdha-1* | Pyruvate dehydrogenase alpha subunit | 0.228 |
| *trx-2* | Thioredoxin | 0.683 |
| *T25B9.9* | Phosphogluconate dehydrogenase | 0.542 |

### *fard-1* and *dhs-25* counteract ferroptosis and aging

To address the potential role of redox-regulated ferroptosis during physiological aging, we then investigated six of the downregulated redox-genes analyzed during normal aging, as possible inhibitors of the ferroptotic process, i.e. *fard-1*, *gpx-7*, *idhb-1*, *ftn-2*, *dhs-25* and *msra-1*. To this extent, available mutant strains were used to evaluate their sensitivity to diethyl maleate (DEM), a GSH-conjugating reagent inducer of ferroptosis, and to quantify lipid peroxidation after acute DEM treatment. Only two of these strains, *fard-1* and *dhs-25* mutants, showed both an increased sensitivity to 10 mM DEM exposure (Figure 4a) and a higher amount of lipid peroxidation (Figure 4b) compared to wild-type worms, confirming their implication in ferroptosis. Intriguingly, *gpx-7* mutant strain exhibited only a significant increase in lipid peroxidation (Figure 4b). Finally, the total glutathione amount as well as the GSH/GSSG ratio were determined at day 0, 4, 7 in all mutant strains. Similar to wild-type, all strains presented with an increased amount of GSH form at day 7 compared to day 0 (Figure S2). Instead, total glutathione content varied significantly in a background and age-dependent manner. Only *ftn-2* mutant displayed no variations in total glutathione amount, while a significant reduction was detected in *gpx-7* mutant at day 4 (Figure 4c). A significant decrease in a time-dependent manner from day 4 was observed in *idhb-1*, *dhs-25* and *msra-1* mutants. Somewhat unexpectedly, total glutathione instead strongly increased from day 4 in *fard-1* mutant. Noteworthy, *fard-1*, *idhb-1*, *ftn-2* and *msra-1* mutant strains presented an amount of glutathione at day 0 completely different from N2: much higher in *idhb-1*, *ftn-2*, *msra-1* and much lower in *fard-1* (Figure 4c).

**Figure 4.**
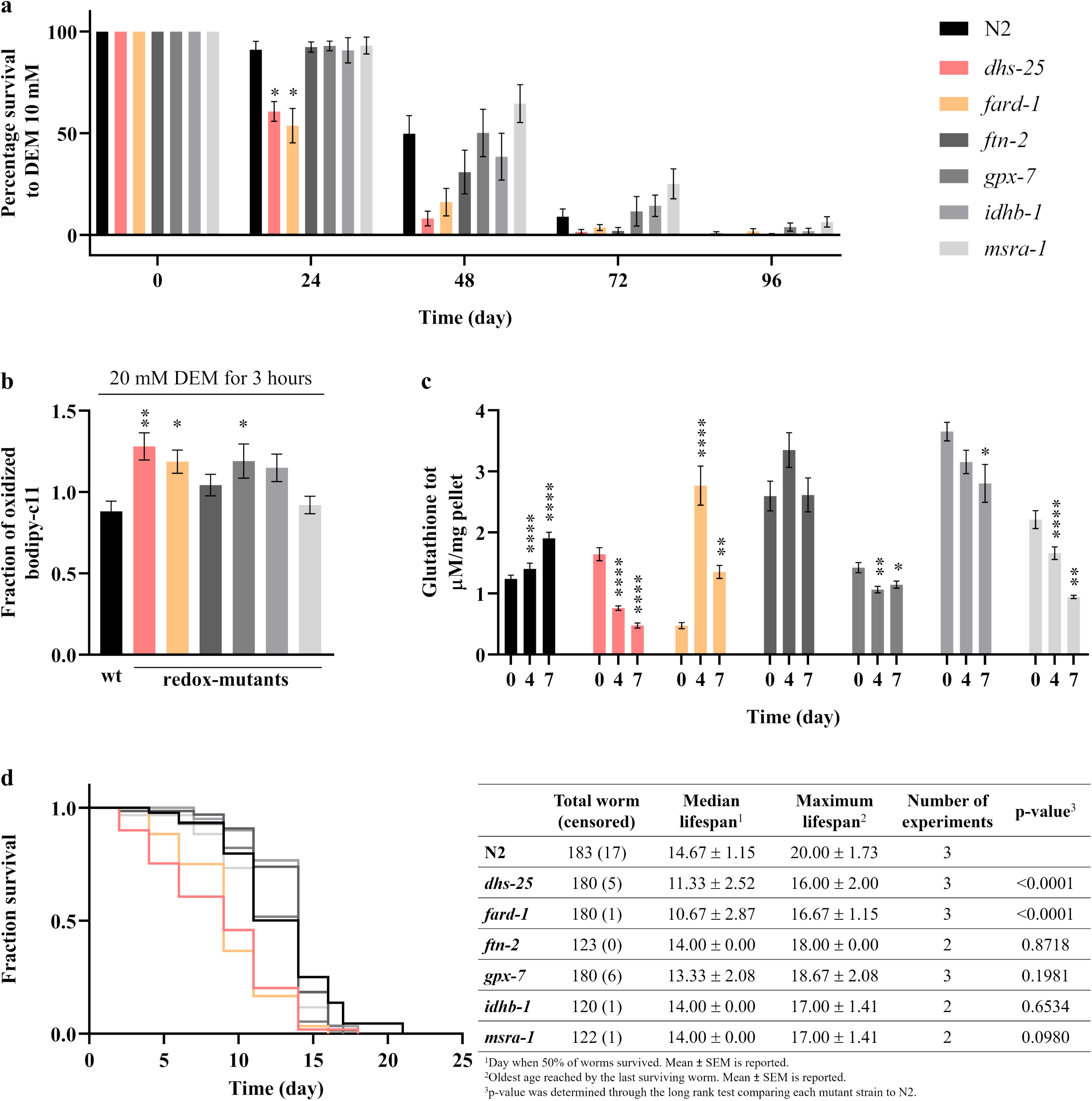
Validation of *fard-1, gpx-7, idhb-1, ftn-2, dhs-25 and msra-1* as inhibitors of ferroptosis, using mutant strains. (a) Survival of 60 N2 and mutant worms to a daily 10 mM DEM exposure was scored for 96 hours. The percentage survival of treated vs untreated animals of each strain was reported as mean ± SEM of at least three biological replicates. (b) Lipid peroxidation was evaluated measuring BODIPY-C11 fluorescence after ferroptosis induction. 2-day adult N2 and mutant strains were treated with 20 mM DEM for 3 hours and then stained with BODIPY-C11. Here, the oxidized/reduced BODIPY-C11 ratio of treated animals normalized on untreated animals for each strain was reported as mean ± SEM of at least three biological replicates and two-way ANOVA with Dunnett’s multiple comparisons test was used for the statistical analysis. (c) Total amount of glutathione in lysates of mutant nematodes at day 0, 4, 7 of adulthood was measured by colorimetric assay. At each specified time point, about 600 nematodes were collected from a population of worms maintained over time. Glutathione amount (μM/mg) was reported as mean ± SEM of at least three biological replicates. *p ≤ 0.05; **p ≤ 0.01; ****p ≤ 0.0001 by one-way ANOVA with Dunnett’s multiple comparisons test. (d) Survival analyses of mutant vs wild-type strains; representative Kaplan-Meier curves were carried out with 60 animals per experiment and lifespan parameters were reported as mean ± SEM of three independent experiments; p-values were calculated through the log rank test.

Next, to further investigate the actual involvement of the selected redox-genes in ferroptosis and aging, lifespan analyses of the mutant strains were carried out. A representative Kaplan-Meier curve for each mutant strain and statistical analysis of the lifespan parameters are reported (Figure 4d). Of note, only *fard-1* and *dhs-25* mutant worms exhibited a significatively shortening of survival compared to N2, displaying a four and three days reduced median lifespan respectively. On the contrary, non-significative differences could be observed between the survival curves of *gpx-7*, *idhb-1*, *ftn-2*, *msra-1* and N2. Nevertheless, all the mutant strains displayed a slightly reduced mean value of maximum lifespan compared to wild-type animals.

Thus *dhs-25* mutant more consistently presents with ferroptosis altered features of relevance for aging. Of note, in support of evolutionarily conserved implication in the ferroptosis process, we found that the expression of its closest mammalian counterpart, the estradiol 17-beta-dehydrogenase 8 (HSD17B8), is significantly reduced in cells which are more sensitive to ferroptosis. We engineered HEK293 cells to carry a point mutation in the frataxin gene (FXNI154F) found in patients with Friedreich’s ataxia, which led to severe depletion of FXN protein expression (Figure 5a, S3). Consistent with the literature, severe frataxin deficiency increased the sensitivity to erastin-induced ferroptosis ^57^, which we found to be prevented by the specific ferroptosis inhibitor Lip-1 (Figure 5b and c). FXN deficient cells also present with significant decreased expression of the HSD17B8 dehydrogenase compared to wild-type cells, while the transcript expression of GPX4 did not change (Figure 5d).

**Figure 5.**
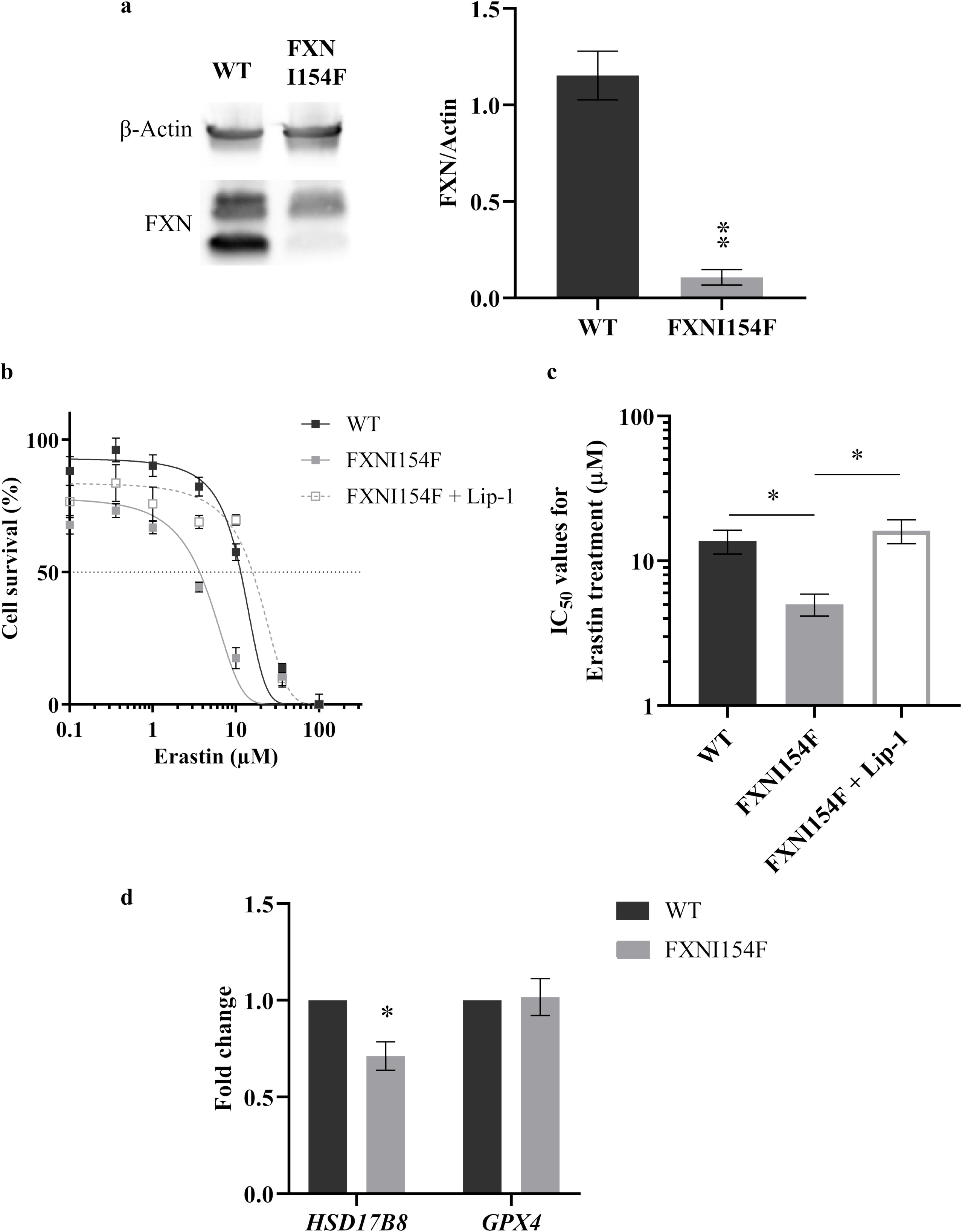
Implications for an evolutionary conserved link between FXN, *dhs-25/HSD17B8* and ferroptosis in mammalian cells. (a) Western blot to analyse frataxin levels in HEK293 cells engineered to carry a point mutation in the frataxin gene (FXNI154F). Left: representative gel, right: quantitative analysis, mean ± SEM of three biological replicates. (b) Survival analyses of both wild-type and mutant HEK293 cells after induction of ferroptosis via increasing concentrations of erastin with or without 100 nM of the ferroptosis inhibitor Liproxstatin-1. Data are shown as mean ± SEM of three biological replicates. (c) IC_50_ values based on (b). (d) Real Time qPCR analyses of HSD17B8 and GPX4 were performed in both HEK293 wild-type and HEK293 FXNI154F. GAPDH was chosen as housekeeping gene for normalization. The fold change value of each gene was reported as mean ± SEM of the biological triplicate. *p ≤ 0.05; **p ≤ 0.01 by unpaired t-test for western blot analysis, one-way ANOVA with Tukey’s multiple comparisons test for cell survival and two-way ANOVA with Bonferroni’s multiple comparisons test for qPCR.

Overall, our study identifies a causal role for ferroptosis in physiological aging in *C. elegans* and indicate NAD-dependent-17-beta-hydroxysteroid dehydrogenase as a novel, evolutionarily conserved gene, whose depletion confers sensitivity to ferroptosis.

## DISCUSSION

Our elderly society has steadily grown in recent years as a direct result of medical and technological advancements. However, the increase in life expectancy is not always accompanied by improved health and well-being ^4^. Therefore, many studies are currently focusing on identifying new strategies to promote healthy aging ^3^. To achieve this, a deeper understanding of the mechanisms underlying the aging process is required. In particular, oxidative damage is a widely recognized hallmark of aging and a progressive general unbalance in redox homeostasis occurs during aging ^8–11^. Oxidative stress has also recently been identified as a key driver of ferroptosis, where especially iron-dependent accumulation of the hydroxyl radical and/or impaired GPX4 activity, lead to PUFA-phospholipid peroxidation and, ultimately, cell death ^18,19^. In the absence of clear evidence of the physiological role of ferroptosis, this work aimed to demonstrate the existence of a strict crosstalk between ferroptosis and aging driven by unbalance in redox homeostasis, using the nematode *C. elegans*. This model organism represents a powerful tool to provide new insights in the aging field thanks to the high conservation of key signaling pathways regulating the aging process ^43,44^. Notably, the first evidences of a link between aging and ferroptosis were indeed derived from studies using *C. elegans* ^37,38^.

At first, the phenotypic characterization of the main healthspan parameters during lifespan confirmed their reliability as indicators of good health: as expected, all analyzed parameters showed age-related deterioration. Nonetheless, aging does not equally affect all physiological parameters and age-associated decline in sensory neurons functionality is for instance much faster than decline in locomotion ^45^. We now find that, while motility and feeding rate progressively decrease from early adulthood (Figure 1b and c), the ability to withstand stressful conditions significantly diminished only in advanced age (Figure 1d and e). According to the literature, age-related decline in musculature is primary due to sarcomere disruption and mainly mitochondrial network breakdown ^58^. Indeed, sarcomeres lose their structural organization and the content of myosin filaments decreases with age ^59^, while muscle mitochondria start to lose their structure already in young adults, leading to reduced ATP production and fragmentation ^60–62^. Muscle deterioration not only contributes to the progressive loss of motility ^63^, but also plays a key role in reduced feeding, while pharyngeal muscles damages are caused by the rapid pumping activity characteristic of the early adulthood ^64^. Moreover, Zhao et al. recently demonstrated that fast pumping in young animals compromises cuticle integrity, facilitating bacterial invasion with the consequent plugging of the pharynx due to the undigested bacteria accumulation ^65,66^. Instead, the ability to withstand external stressors (i.e. high temperatures and hydrogen peroxide exposure) that affects proteostasis, redox balance, metabolism and membrane integrity ^48^, declines starting from one-week old nematodes, suggesting a slower physiological deterioration of the cellular pathways responsible for homeostasis maintenance (Figure 1d and e). On day 4, thermal resistance remains unchanged with respect to day 0 and a slight non-significative increment in hydroxyl radical level was observed. Moreover, the antioxidant defenses seem to be sufficient to maintain a balanced GSH/GSSG ratio. Differently, redox homeostasis begins to be challenged starting from day 7, when a slightly diminished ability to counteract thermal stress was observed, with a significative increase in Fenton reaction-produced hydroxyl radical and a necessary synthesis of new GSH to mitigate the oxidized environment. However, only at day 14 the worms exhibit both a dramatic increment in hydroxyl radical formation and a drop in total glutathione, leading to a drastic loss of the capacity to counter external stressors. At the median lifespan, animals quite completely lose their redox homeostasis even though GSSG continues to be reduced, and this could be correlated to the higher amount of ongoing Fenton reaction and the impaired synthesis of new glutathione. The limited amount of antioxidant reserve is not sufficient to maintain homeostasis, even though GSSG continues to be recycled into the scavenger GSH form (Figure 2). In agreement with this, an upregulation of *gcs-1*, the gene encoding glutamate-cysteine ligase responsible for the first step of glutathione biosynthesis, has been observed between day 4 and 7, followed by a progressive downregulation from day 7 and to day 14 ^37^. Moreover, our qPCR data showed a downregulation of specific genes with oxidoreductase activity during wild-type aging, supporting that a reduced expression of these genes contributes to the breakdown of cellular redox homeostasis (Figure 3). These genes were selected because they were differentially expressed upon mild frataxin depletion, a condition that hormetically extends lifespan in *C. elegans,* at least in part through the inhibition of ferroptosis ^37^. The upregulation of *fard-1*, *gpx-7*, *idhb-1*, *ftn-2*, *dhs-25* and *msra-1* in *frh-1* RNAi worms suggests that these genes may function as inhibitors of ferroptosis and their observed downregulation during wild-type aging is consistent with the hypothesis that ferroptosis increases with age. Among these, the fatty acyl-CoA reductase *fard-1*, the glutathione peroxidase *gpx-7* and the iron storage protein ferritin *ftn-2* have been previously implicated in *C. elegans* aging and their involvement in ferroptosis is inferred based on their mammalian orthologs ^22,67,68^. Instead, the dehydrogenases *idhb-1* and *dhs-25* and the reductase responsible for detoxification of methionine sulfoxides, *msra-1*, emerge as novel and interesting candidates for further investigation of their role in ferroptosis. Our results demonstrated that only *fard-1* and *dhs-25* mutant animals, display higher sensitivity to the ferroptosis inducer DEM, increased lipid peroxidation, anticipated drop in total glutathione and reduced lifespan, compared to wild-type, confirming their involvement in ferroptosis-aging association (Figure 4 and 5). In contrast, the *gpx-7* mutants displayed altered lipid peroxidation and total glutathione levels, although neither alteration in the DEM response nor in the lifespan could be detected (Figure 4). A possible explanation for these somewhat unexpected results lies in the fact that *C. elegans* possesses four isoforms, *gpx-1*, *gpx-2*, *gpx-6*, *gpx-7*, all of them encoding for the mammalian ortholog GPX4, the sole enzyme responsible for the detoxification of PUFA-phospholipid hydroperoxide ^69^. As a result, in slower processes like resistance to DEM and lifespan, isoforms might compensate for the lack of *gpx-7*, while this is not occurring in early or acute processes, such as lipid peroxidation and glutathione depletion. Noteworthy, *fard-1* is one of the genes involved in the synthesis of ether lipids, whose beneficial role in aging and longevity has been recently recognized ^70^. Loss of function mutations in one of the enzymes responsible for ether lipids biosynthesis, including *fard-1*, lead to a reduced production of ether-bound lipids and to a shortened lifespan in *C. elegans* ^71,72^. Furthermore, Cedillo et al. demonstrated that the single overexpression of *fard-1* is sufficient to extend lifespan in worms in a *skn-1*, *aak-2* and *daf-16* dependent way ^73^. On the other hand, ether lipids can extend lifespan by modulating ferroptosis: unlike in mammalian cells, these species are more likely to be contained in monounsaturated (MUFA) rather than polyunsaturated fatty acids (PUFA) in *C. elegans* ^67,74,75^. High levels of MUFAs can displace PUFAs from membrane phospholipids, thus protecting against peroxidation ^76^. Instead, *dhs-25*, ortholog of the mammalian hydroxysteroid 17-beta dehydrogenase 8, encodes for a NAD-dependent short chain dehydrogenase potentially responsible for mitochondrial fatty acid synthesis ^77^, but still poorly characterized and so far never implicated in *C. elegans* aging. Moreover, to the best of our knowledge, no evidence of HSD17B8 implication in ferroptosis is available neither in *C. elegans* nor in mammals. Nonetheless, Wei et al. recently showed that a low expression of the ferroptosis-related gene HSD17B11 (member of the same group of alcohol dehydrogenases HSD17B) displays increased tumor recurrence and poor prognosis in patients affected by lung cancer ^78^. In addition, the dehydrogenase activity of *dhs-25* can be both NAD^+^ and NADP^+^ dependent ^79^, so its role in ferroptosis regulation could be directly related to its involvement in the GSSG recycling process, or in lipid metabolism. In conclusion, our work demonstrates the existence of a causal connection between ferroptosis and physiological aging triggered by age-dependent alteration in redox homeostasis and identified novel genes involved in both processes. Future studies are needed to further elucidate the roles of *fard-1* and *dhs-25* in the ferroptosis-aging connection not only in the *C. elegans* model but also in mammalian systems.

## MATERIALS AND METHODS

### C. elegans

#### *C. elegans* strains, maintenance conditions and synchronization

The following strains were used in this work: wild-type N2 Bristol, BX275: *fard-1(wa28) X*., RB1830: *idhb-1(ok2368) X*., RB668: *ftn-2(ok404) I*., FX02166: *gpx-7(tm2166) X*., FX04291: *dhs-25(tm4291) X*., FX01421: *msra-1(tm1421) II*. N2, BX275, RB1830 and RB668 were provided by the Caenorhabditis Genetics Center (CGC), University of Minnesota. Whereas FX02166, FX04291, FX01421 come from the National BioResource Project (*C. elegans*), Tokyo Women’s Medical University School of Medicine. All the strains were maintained at 20 °C on Nematode Growth Medium plates (NGM; 2.5 g/L bacteriological peptone, 3 g/L NaCl, 17 g/L agar, 1 mM CaCl_2_, 1 mM MgSO_4_, 1 mM cholesterol, 25 mM KH_2_PO_4_) using alive *Escherichia coli* OP50 strain at the OD_600_=0.3 as food source. Egg lay was carried out to synchronize worms in all the experiments. Briefly, gravid adult worms were allowed to lay eggs for 16 hours and then removed. Newly laid eggs were grown for 3 days at 20 °C, until reaching the first day of adulthood (day 0). All the experiments were conducted using live OP50 at OD_600_=0.3 at 20 °C, unless specified.

#### Lifespan analysis

A population of 60 animals at the first day of adulthood was scored for survival analysis. 40 μM 5-Fluoro-2-deoxyuridine (FuDR, Invitrogen Corporation, Massachusetts, USA) was added to NGM plates during the fertile period to avoid eggs hatching (first week). Living worms were counted and transferred every other day until the last surviving animal. Animals unable to react after a platinum wire touch and with no more pharyngeal pumping were scored as dead, while nematodes were censored if missing or having crawled off the plate.

#### Motility assay

Worms’ locomotion was evaluated by counting the number of body bends per minute. Every time the head of the worm drew a curve to the right or to the left in the direction of movement, one body bend was counted ^47^. A starting population of 200 nematodes at day 0 was scored every day during the first week of adulthood and every other day in the following weeks, at least in biological triplicate. The collected data were represented as box plots of the body bends/min counted at each time of life analyzed.

#### Pharyngeal pumping rate

Pumping rate assay was used to measure the feeding rate of the animals ^80^. 30 nematodes were scored for their pharyngeal pulses every other day during lifespan, until day 16 of adulthood. Since the pharynx contractions are too fast to be counted, the animals were recorded using a SteREO Discovery V12 microscope (Carl Zeiss Microscopy GmbH, Germany) and movies were replied at a half of their speed. Data were collected in biological triplicate and represented as box plots.

#### Thermal stress resistance assay

Nematode survival at the non-permissive temperature of 37 °C was measured to assess *C. elegans* ability to contrast heat stress. NGM plates were pre-heated at 37 °C for 20 minutes before laying the animals and starting the experiment. 30 synchronous nematodes at day 0 were scored for survival at 37 °C every two days of their lifespan, until day 16 of adulthood.

#### Oxidative stress resistance assay

Nematode survival in the presence of hydrogen peroxide was measured to assess *C. elegans* ability to contrast oxidative stress. NGM plates containing 1 μL/mL of H_2_O_2_ 30% w/w (Sigma-Aldrich/Merck, Darmstadt, Germany) and seeded with *E. coli* OP50 OD_600_=0.3 were freshly prepared and kept in the dark before and during use. 30 synchronized animals were scored for survival on H_2_O_2_ plates every two days of their lifespan, until day 16 of adulthood.

#### Reactive Oxygen Species measurement

To assess the redox state of the worms during aging, Fenton reaction was measured at day 0, 4, 7 and 14 of adulthood. 100 nematodes for each time point were collected in phosphate buffered saline buffer (PBS; 20 mM KH_2_PO_4_, 20 mM K_2_HPO_4_, 150 mM NaCl; pH 7.2), washed four times and centrifuged at 2 000 xg, 4 °C for 1 minute. The nematodes pellet was then snapped frozen in liquid nitrogen and kept at −80 °C until use. Once an entire biological set had been collected, worms were lysed through homogenization (four cycles of 30 seconds homogenization + 30 seconds in ice), followed by sonication (five cycles of 1 second at 10% amplitude with intervals of 30 seconds in ice). Protein amount was determined through the Bradford protein assay ^81^ and 10 μg of the lysate were then incubated with 50 μM 2’,7’-dichlorodihydrofluorescein-diacetate (H_2_DCFDA, Sigma-Aldrich/Merck, Darmstadt, Germany) at 37 °C for 1 hour. This probe allows to estimate H_2_O_2_- and iron-dependent formation of the hydroxyl radical ^82^. DCF fluorescence was measured through a multiplate reader (Victor 3, PerkinElmer, Waltham, MA, USA) at the excitation/emission wavelengths of 485 and 520 nm, respectively.

#### Total glutathione and GSH/GSSG ratio measurement

Total glutathione amount and the distribution of its reduced (GSH) and oxidized (GSSG) forms were measured at day 0, 4, 7 and 14 of adulthood. Around 400-600 nematodes at each time point were washed in PBS buffer, centrifuged at 2 000 xg, 4°C for 1 minute and the resulting pellet was weighted, snap frozen in liquid nitrogen and stored at −80 °C. Once an entire biological set had been collected, worms were lysed through homogenization (three cycles of 30 seconds homogenization + 30 seconds ice) in 250 μL of a 1% sulfo-salicylic acid solution. To quantify the total glutathione amount, a 96-well plate was filled as follows: 5 μL of the worm lysates, 95 μL of MilliQ water, 100 μL of reaction mixture (100 μM DTNB (5,5′-dithiobis(2-nitrobenzoic acid)), 200 μM NADPH and 0.46 U/mL glutathione reductase in a 100 mM sodium phosphate buffer pH 7.5). A calibration curve with a GSH standard (0-10 μM) was prepared and processed like the samples. For GSSG quantification, 100 μL of the worm lysates were treated with 2 μL of 2-vinylpyridine solution (27 μL 2-vinylpyridine in 98 μL ethanol) for 1 hour at room temperature. 2-vinylpyridine masks SH-groups, allowing to quantify only GSSG. A 96-well plate was then filled as follows: 5 μL of the worm lysates, 95 μL of MilliQ water, 100 μL of reaction mixture (as described before). A calibration curve with a GSSG standard (0-10 μM) was prepared and processed like the samples. For both the enzymatic assays, absorbance at 405 nm was measured immediately and after 50 minutes of incubation at room temperature using a multiplate reader (SPECTROstarNano, BMG Labtech, Ortenberg, Germany). Total glutathione and GSSG were determined based on their standard curves and normalized on pellet weight, expressed in mg. Subtracting the amount of GSSG from the total glutathione quantity, the level of GSH was calculated. The ratio GSH/GSSG was then expressed as a percentage value for each time point of lifespan.

### Genes expression

#### RNA extraction and reverse transcription

RNA was extracted from wild type animals at day 0, 4, 7 and 14 of adulthood fed OP50 or from animals fed for three consecutive generation with HT115 bacteria transformed either with empty vector (pL4440) or with vector expressing *frh-1* dsRNA. Around 1 000-2 000 animals for each time point were collected in PBS buffer, washed four times and centrifuged at 2 000 xg, 4 °C for 1 minute. The pellet was then snap frozen in liquid nitrogen and stored at −80 °C. Once an entire biological set had been collected, RNA was extracted according to the RNeasy Plus Mini kit (QIAGEN, Hilden, Germany) procedure. Briefly, the highly denaturing guanidine isothiocyanate–containing buffer was added to the samples and worms were sonicated in ice performing 2 cycles of 10 seconds at 30% intensity and 1 cycle of 10 seconds at 40% intensity, with 1 minute pause between each cycle. The homogenized lysates were treated to get rid of gDNA and pure RNA was obtained. 0.1 μg/μL of the resulting RNA was heated to 60 °C for 5 minutes and cooled on ice for another 5 minutes to release the possible secondary structures present. Then, RNA was reverse transcribed into cDNA following the procedure of the GoScript^TM^ Reverse Transcription Mix, Oligo(dT) protocol (Promega, Madison, USA). Reaction products were stored at −20 °C. RNA from at least three biological replicates was extracted and reverse transcribed.

#### Quantitative Real Time PCR

cDNAs obtained at the different time life were used for the qPCR assay. Each reaction mixture, in triplicate, was prepared as follows: iQTM SYBR^®^ Green Supermix (BIO-RAD, California, USA) 1x, 300 nM forward primer, 300 nM reverse primer, 10 ng cDNA template, DNAse free water to the final volume of 20 μL (values of final concentrations are indicated). PCR was performed according to the default thermal protocol of the QuantStudioTM 3 Real-Time PCR System machine (ThermoFisher Scientific, Massachusetts, USA):

- Hold stage: Step 1, 50 °C – 2 minutes; Step 2, 95 °C – 10 minutes.
- PCR stage: Step1, 95 °C – 15 seconds; Step 2, 60 °C – 1 minute. x40 cycles.
- Melt curve stage: Step 1, 95 °C – 15 seconds; Step 2, 60 °C – 1 minute; Step 3 (dissociation), 95 °C – 1 second.

The primers were designed using Primer BLAST tool (NCBI, Maryland, USA), purchased by Sigma-Aldrich/Merck (Darmstadt, Germany) and their efficiency and specificity was verified in cDNA samples of worms at day 0 (Table S2). *cdc-42* was selected as the housekeeping gene. Real time qPCRs were performed in biological triplicate and data were analysed based on the 2^-ΔΔCt^ method.

#### Diethyl maleate sensitivity test

Diethyl maleate (DEM, Sigma-Aldrich/Merck, Darmstadt, Germany) was directly added to NGM at the final concentration of 10 mM. 60 animals at day 0 of adulthood were scored for survival in the presence of DEM every 24 hours for five days. Data were expressed as percentage of survival and reported as the ratio between treated and untreated animals at each time point.

#### Lipid peroxidation measurement

The level of lipid peroxidation after ferroptosis induction was determined using the BODIPY-C11 probe (no. 27086, Caymann chemical, Michigan, USA). This probe was designed to be able to reach the lipid bilayer and to shift its fluorescent emission from red to green upon oxidation by phospholipids peroxyl radicals, thus reflecting the level of lipid peroxidation in a sample ^83^. 70 2-day adult worms were treated with 20 mM DEM on plate for 3 hours, washed three times with M9 buffer (22 mM KH_2_PO_4_, 42 mM Na_2_HPO_4_, 86 mM NaCl, 1 mM MgSO_4_) and stained with 500 μL BODIPY-C11 solution (M9, 10 μM BODIPY-C11 and 67 μL/mL of OP50 OD_600_=0.3) for 45 minutes. Additional untreated and unstained worms were also processed. Then, worms were washed three times in M9 and 125 μL of each were aliquot in black 96-well plates to measure fluorescence. A TECAN (Infinite M Nano+, Männedorf, Switzerland) multiplate reader was used. Oxidized BODIPY-C11 was acquired using 488 excitation and 530 emission wavelengths, while reduced BODIPY-C11 was measured using 543 excitation and 590 emission wavelengths. Background fluorescence of unstained worms was subtracted and the ratio between oxidized and reduced BODIPY-C11 was calculated for each treated and untreated sample. Finally, the oxidized/reduced BODIPY-C11 ratio of treated worms was normalized on the untreated animals.

### Mammalian Cells

#### Cell culture

Human embryonic kidney (HEK) cells were cultured in DMEM medium, containing 4.5 g/L glucose, 10 % FCS and 1 % penicillin/streptomycin, and cultivated at 37 °C in a humidified atmosphere containing 5% CO_2_.

#### Cell survival

The cell survival was measured in a 96 well plate containing 1×10^4^ cells per well. The cells were treated with increasing concentrations of erastin (Sigma-Aldrich, USA) for 24 h to induce ferroptosis and co-treated with or without 100 nM Liproxstatin-1 (Sigma-Aldrich, USA) to inhibit ferroptosis. Afterwards cells were incubated with CellTiter-Blue® reagent (Promega, USA) for 1 h at 37 °C and 5 % CO_2_. Fluorescence was measured with TECAN plate reader with an excitation at 562 nm and an emission at 590 nm.

#### Protein quantification

The protein level of supernatants of lysed cells were measured by using the BC assay quantification kit (Interchim, France) according to the manufacturer’s protocol.

#### Electrophoresis and western blot

Samples were diluted in 1x Laemmli protein sample buffer (Bio-Rad, USA) and incubated for 5 min at 95 °C. Proteins were separated by mass in the Mini-PROTEAN® TGX™ Precast Gel (Bio-Rad, USA) in the Mini-PROTEAN® Tetra System (Bio-Rad, USA) with running buffer *(25 mM Tris, 192 mM Glycine, 0.1% SDS, pH 8.3)* for approximately 1 h at 130 V. To detect specific proteins the gel was blotted on a 0.2 µM nitrocellulose membrane using the Trans-Blot Turbo Transfer System (Bio-Rad) for 7 min at 1.3 A and 25 V. The membrane was blocked with blocking buffer (PBS, 5 % BSA, 0.05 % tween) for 1 h and afterwards stained with primary antibodies against Frataxin and ß-Actin (1:1000 in PBS, 2.5 % BSA, 0.05 % tween) at 4 °C over night. After 3 washing steps with PBS containing 0.05 % tween, membranes were incubated with fluorescent secondary antibodies (1:15000 in PBS, 2.5 % BSA, 0.05 % tween) for 1 h at RT. The detection was performed using the ChemiDoc Imaging System (Bio-Rad, USA).

#### qPCR

RNA was isolated from cells using the TRIzol Reagent (Invitrogen, USA) according to manufacturer’s protocol. 1.5 µg was transcripted to cDNA using Reverse Transcription Kit (Applied Biosystems, USA). Quantitative PCRs were performed in technical triplicates and biological triplicates using PowerUp SYBR Green Master Mix and QuantStudio 3 (Applied Biosystems, USA). Mean values were normalized to the housekeeping gene GAPDH (Qiagen, Germany) and fold gene expression was calculated in relation to wild-type HEK cells.

Primer sequences:

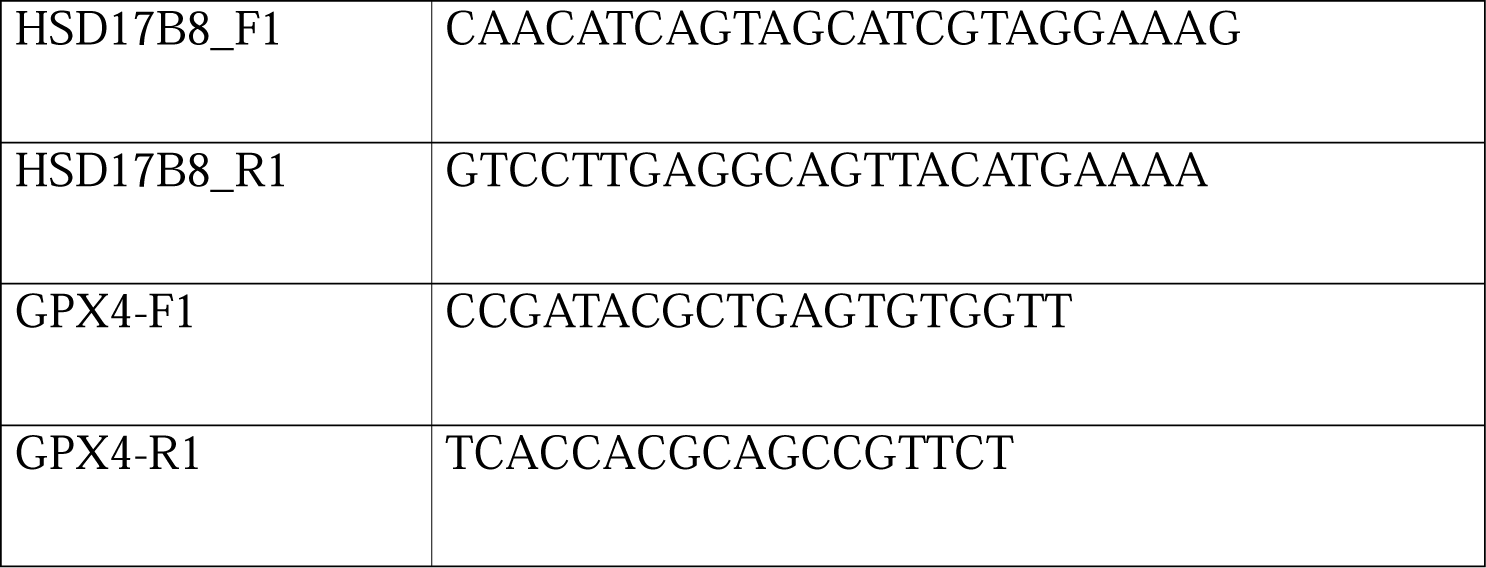

#### Quantification and Statistical analysis

GraphPad Prism 8.0.2 software was used to analyse all data. Survival curves were represented as Kaplan-Meier and p-values calculated through the log rank test. Comparison between groups were performed using unpaired t-test; one-way ANOVA with Dunnett’s or Tukey’s multiple comparisons test; Kruskal-Wallis, Dunn’s multiple comparisons test; two-way ANOVA with Dunnett’s or Bonferroni’s multiple comparisons test. Collected data were shown as mean ± SEM of at least three independent biological replicates. Statistical relevance was determined according to the following p-values: * p < 0.05, ** p < 0.01, *** p < 0.001, **** p < 0.001.

**Supplementary information is available at *Cell Death & Differentiation*’s website.**

## ACKNOWLEDGEMENTS

We would like to thank the Caenorhabditis Genetics Center (funded by the National Institutes of Health Office of Research Infrastructure Programs: P40OD010440), as well as the National Bioresource Project (NBRP) for *C. elegans* strains.

## AUTHOR CONTRIBUTION STATEMENT

Study concept and design: RP, MF, PF, CB, NV, and MER. Development of methodology and writing, review and revision of the paper: RP, BS, FB, MF, PF, CB, NV and MER. Acquisition, analysis and interpretation of data, and statistical analysis: RP, BS, FB, MF, LS, SM and LT. All authors read and approved the final paper.

## DATA AVAILABILITY STATEMENT

The original contributions presented in the study are included in the article and supplementary material, further inquiries can be directed to the corresponding authors.

