## Supplementary material for "Redox homeostasis in ferroptosis and aging: a causal role for *fard-1* and *dhs-25* in *Caenorhabditis elegans*": Supplementary information.docx

**Table S1**. Description and corresponding mammalian ortholog of the genes chosen for gene expression analysis during lifespan (Separate excel file).

**Table S2**. List of forward and reverse primers used in quantitative real time PCR assays (*C. elegans* experiments; Separate excel file).

**Figure S1.** Validation of selected redox-genes expression in frataxin silenced animals. Real Time PCR analysis of the oxdoreductase genes selected from frh-1 RNAi worms trascriptomic analysis were carried out. Around 1 000-2 000 N2 and frh-1 RNAi nematodes were collected for RNA extraction and reverse transcription. The fold change value of each gene was reported as mean ± SEM of a biological triplicate. *p ≤ 0.05; **p ≤ 0.01; ***p ≤ 0.001; ****p ≤ 0.0001 by Kolmogorov-Smirnov test in GraphPad Prism 8.0.2.

**Figure S2**. Evaluation of the ratio between reduced/oxidized glutathione in lysates of *dhs-25* (a), *fard-1* (b), *ftn-2* (c), *gpx-7* (d), *idhb-1* (e) and *msra-1* (f) mutant nematodes at day 0, 4 and 7 of adulthood. A colorimetric assay exploiting 5,5′- dithiobis (2-nitrobenzoic acid) reduction was performed to determine the glutathione total amount and the level of the oxidized form (GSSG). Levels of the reduced form (GSH) were derived by subtraction. A population of each mutant strain was maintained at 20 °C by transferring the animals on fresh NGM plates seeded with *E. coli* OP50 OD_600_=0.3 every day. At each specified time point, about 600 nematodes were collected and frozen. Lysates were processed as described in par. “Total glutathione and GSH/GSSG ratio measurement” (Met&Mat) to quantify total glutathione and GSSG, after the incubation with the GSH masking reagent 2-vinylpyridine. Percentage values of GSH and GSSG respect to total glutathione for each time point were represented as mean ± SEM of at least three biological replicates. *p ≤ 0.05; **p ≤ 0.01; ***p ≤ 0.001 by two-way ANOVA with Dunnett’s multiple comparisons test.

**Figure S3**. Western blot to analyze frataxin levels in HEK293 cells engineered to carry a point mutation in the frataxin gene (FXNI154F). Representative gel of the three biological replicates (a-c).
