## Supplementary material for "Redox homeostasis in ferroptosis and aging: a causal role for *fard-1* and *dhs-25* in *Caenorhabditis elegans*": Table S1.pdf

| Gene | WormBase ID | Protein | Ortholog name | ENSG_Ortholog | Description |
|---|---|---|---|---|---|
| *acdh-2* | WBGene00015894 | Acyl CoA dehydrogenase | ACAD10 | ENSG00000111271 | acyl-CoA dehydrogenase family member 10 [Source:HGNC Symbol;Acc:HGNC:21597] |
| | | | ACADL | ENSG00000115361 | acyl-CoA dehydrogenase long chain [Source:HGNC Symbol;Acc:HGNC:88] |
| | | | ACADM | ENSG00000117054 | acyl-CoA dehydrogenase medium chain [Source:HGNC Symbol;Acc:HGNC:89] |
| | | | ACADS | ENSG00000122971 | acyl-CoA dehydrogenase short chain [Source:HGNC Symbol;Acc:HGNC:90] |
| | | | ACOXL | ENSG00000153093 | acyl-CoA oxidase like [Source:HGNC Symbol;Acc:HGNC:25621] |
| | | | ACOX1 | ENSG00000161533 | acyl-CoA oxidase 1 [Source:HGNC Symbol;Acc:HGNC:119] |
| | | | ACOX2 | ENSG00000168306 | acyl-CoA oxidase 2 [Source:HGNC Symbol;Acc:HGNC:120] |
| | | | ACAD9 | ENSG00000177646 | acyl-CoA dehydrogenase family member 9 [Source:HGNC Symbol;Acc:HGNC:21497] |
| | | | ACADSB | ENSG00000196177 | acyl-CoA dehydrogenase short/branched chain [Source:HGNC Symbol;Acc:HGNC:91] |
| *acdh-9* | WBGene00017874 | Acyl CoA dehydrogenase | ACAD8 | ENSG00000151498 | acyl-CoA dehydrogenase family member 8 [Source:HGNC Symbol;Acc:HGNC:87] |
| *cdc-42* | WBGene00000390 | Cell division cycle related | CDC42 | ENSG00000070831 | cell division cycle 42 [Source:HGNC Symbol;Acc:HGNC:1736] |
| *ctl-3* | WBGene00013220 | Catalase | CAT | ENSG00000121691 | catalase [Source:HGNC Symbol;Acc:HGNC:1516] |
| *cyp-13A8* | WBGene00011674 | Cytochrome P450 family | N.A. | N.A. | N.A. |
| *cyp-14A1* | WBGene00010705 | Cytochrome P450 family | N.A. | N.A. | N.A. |
| *cyp-14A4* | WBGene00011009 | Cytochrome P450 family | N.A. | N.A. | N.A. |
| *cyp-33C8* | WBGene00019967 | Cytochrome P450 family | N.A. | N.A. | N.A. |
| *dhs-18* | WBGene00000981 | Dehydrogenases, short chain | RDH8 | ENSG00000080511 | retinol dehydrogenase 8 [Source:HGNC Symbol;Acc:HGNC:14423] |
| | | | DHRS7 | ENSG00000100612 | dehydrogenase/reductase 7 [Source:HGNC Symbol;Acc:HGNC:21524] |
| | | | DHRS2 | ENSG00000100867 | dehydrogenase/reductase 2 [Source:HGNC Symbol;Acc:HGNC:18349] |
| | | | DHRS7B | ENSG00000109016 | dehydrogenase/reductase 7B [Source:HGNC Symbol;Acc:HGNC:24547] |
| | | | HSD11B1 | ENSG00000117594 | hydroxysteroid 11-beta dehydrogenase 1 [Source:HGNC Symbol;Acc:HGNC:5208] |
| | | | HSDL2 | ENSG00000119471 | hydroxysteroid dehydrogenase like 2 [Source:HGNC Symbol;Acc:HGNC:18572] |
| | | | DHRS4 | ENSG00000157326 | dehydrogenase/reductase 4 [Source:HGNC Symbol;Acc:HGNC:16985] |
| | | | CBR1 | ENSG00000159228 | carbonyl reductase 1 [Source:HGNC Symbol;Acc:HGNC:1548] |
| | | | CBR3 | ENSG00000159231 | carbonyl reductase 3 [Source:HGNC Symbol;Acc:HGNC:1549] |
| | | | HSD11B1L | ENSG00000167733 | hydroxysteroid 11-beta dehydrogenase 1 like [Source:HGNC Symbol;Acc:HGNC:30419] |
| | | | DHRS7C | ENSG00000184544 | dehydrogenase/reductase 7C [Source:HGNC Symbol;Acc:HGNC:32423] |
| | | | DHRS4L2 | ENSG00000187630 | dehydrogenase/reductase 4 like 2 [Source:HGNC Symbol;Acc:HGNC:19731] |
| | | | DHRS11 | ENSG00000278535 | dehydrogenase/reductase 11 [Source:HGNC Symbol;Acc:HGNC:28639] |
| *dhs-25* | WBGene00000988 | Dehydrogenases, short chain | HSD17B8 | ENSG00000112474 | hydroxysteroid 17-beta dehydrogenase 8 [Source:HGNC Symbol;Acc:HGNC:3554] |
| *dld-1* | WBGene00010794 | Dihydrolipoamide dehydrogenase | DLD | ENSG00000091140 | dihydrolipoamide dehydrogenase [Source:HGNC Symbol;Acc:HGNC:2898] |
| *fard-1* | WBGene00022200 | Fatty acyl-CoA reductase | FAR2 | ENSG00000064763 | fatty acyl-CoA reductase 2 [Source:HGNC Symbol;Acc:HGNC:25531] |
| | | | FAR1 | ENSG00000197601 | fatty acyl-CoA reductase 1 [Source:HGNC Symbol;Acc:HGNC:26222] |
| *fat-6* | WBGene00001398 | Fatty acid desaturase | SCD | ENSG00000099194 | stearoyl-CoA desaturase [Source:HGNC Symbol;Acc:HGNC:10571] |
| | | | SCD5 | ENSG00000145284 | stearoyl-CoA desaturase 5 [Source:HGNC Symbol;Acc:HGNC:21088] |
| *ftn-2* | WBGene00001501 | Ferritin | FTL | ENSG00000087086 | ferritin light chain [Source:HGNC Symbol;Acc:HGNC:3999] |
| | | | FTHL17 | ENSG00000132446 | ferritin heavy chain like 17 [Source:HGNC Symbol;Acc:HGNC:3987] |
| | | | FTH1 | ENSG00000167996 | ferritin heavy chain 1 [Source:HGNC Symbol;Acc:HGNC:3976] |
| | | | FTMT | ENSG00000181867 | ferritin mitochondrial [Source:HGNC Symbol;Acc:HGNC:17345] |
| *gdh-1* | WBGene00014095 | Glutamate dehydrogenase | GLUD1 | ENSG00000148672 | glutamate dehydrogenase 1 [Source:HGNC Symbol;Acc:HGNC:4335] |
| | | | GLUD2 | ENSG00000182890 | glutamate dehydrogenase 2 [Source:HGNC Symbol;Acc:HGNC:4336] |
| *gpx-7* | WBGene00019846 | Glutathione peroxidase | GPX4 | ENSG00000167468 | glutathione peroxidase 4 [Source:HGNC Symbol;Acc:HGNC:4556] |
| *idh-1* | WBGene00010317 | Isocitrate dehydrogenase | IDH1 | ENSG00000138413 | isocitrate dehydrogenase (NADP(+)) 1 [Source:HGNC Symbol;Acc:HGNC:5382] |
| *idhb-1* | WBGene00007993 | Isocitrate dehydrogenase beta | IDH3B | ENSG00000101365 | isocitrate dehydrogenase (NAD(+)) 3 non-catalytic subunit beta [Source:HGNC Symbol;Acc:HGNC:5385 |
| *msra-1* | WBGene00018393 | Methionine sulfoxide reductase A | MSRB2 | ENSG00000148450 | methionine sulfoxide reductase B2 [Source:HGNC Symbol;Acc:HGNC:17061] |
| | | | MSRA | ENSG00000175806 | methionine sulfoxide reductase A [Source:HGNC Symbol;Acc:HGNC:7377] |
| | | | MSRB1 | ENSG00000198736 | methionine sulfoxide reductase B1 [Source:HGNC Symbol;Acc:HGNC:14133] |
| *pdha-1* | WBGene00011510 | Pyruvate dehydrogenase alpha subunit | PDHA1 | ENSG00000131828 | pyruvate dehydrogenase E1 subunit alpha 1 [Source:HGNC Symbol;Acc:HGNC:8806] |
| | | | PDHA2 | ENSG00000163114 | pyruvate dehydrogenase E1 subunit alpha 2 [Source:HGNC Symbol;Acc:HGNC:8807] |
| *trx-2* | WBGene00007099 | Thioredoxin | TXN2 | ENSG00000100348 | thioredoxin 2 [Source:HGNC Symbol;Acc:HGNC:17772] |
| *T25B9.9* | WBGene00012015 | Phosphogluconate dehydrogenase | PGD | ENSG00000142657 | phosphogluconate dehydrogenase [Source:HGNC Symbol;Acc:HGNC:8891] |
