## Supplementary material for "Redox homeostasis in ferroptosis and aging: a causal role for *fard-1* and *dhs-25* in *Caenorhabditis elegans*": Table S2.pdf

| Gene | WormBase ID | Primer Name | Primer Sequence 5'->3' | Amplicon Size |
|---|---|---|---|---|
| *acdh-2* | WBGene00015894 | ACDH2_FOR<br>ACDH2_REV | CCGAAGTATGGAGGATCTGGAG<br>CGAAGGCGCCAACCAAATTC | 149 |
| *acdh-9* | WBGene00017874 | ACDH9_FOR<br>ACDH9_REV | GCAGATTGGGATAAGCAGGGAG<br>TATAAGCTGCCGTCGACACG | 175 |
| *cdc-42* | WBGene00000390 | CDC42_FOR<br>CDC42_REV | ATTACGCCGTCACAGTAATG<br>ATCCCTGAGATCGACTTGAG | 248 |
| *ctl-3* | WBGene00013220 | CTL3_FOR<br>CTL3_REV | TCCCCACATGGTCAATCTAACG<br>GCAGGTGGGGTTCCTGATTTC | 101 |
| *cyp-13a8* | WBGene00011674 | CYP13A8_FOR<br>CYP13A8_REV | AGCTCAAGGTTTTAGATGGA<br>CATAAGCTCCTGTGTGCTAT | 112 |
| *cyp-14a1* | WBGene00010705 | CYP14A1_FOR<br>CYP14A1_REV | TTCACCAGTTCCCTCCAGAC<br>AGAACGGTGAGCTCGGTAAG | 176 |
| *cyp-14a4* | WBGene00011009 | CYP14A4_FOR<br>CYP14A4_REV | TTGGATGTTGTTGGCAACTC<br>TTGGTATCCTTATTGCTTTG | 143 |
| *cyp-33c8* | WBGene00019967 | CYP33C8_FOR<br>CYP33C8_REV | GATGATGTGCTCAACTACTG<br>CTTGAGCCTTTTGCTTCTTC | 151 |
| *dhs-18* | WBGene00000981 | DHS18_FOR<br>DHS18_REV | AGCAGCGAAGACAGCAACTG<br>TACTGCTGCCTCATCACGAAC | 127 |
| *dhs-25* | WBGene00000988 | DHS25_FOR<br>DHS25_REV | CCGGAGTCTTCCATGTATCCC<br>GTGGCTGCGTAGTTGGTTTG | 133 |
| *dld-1* | WBGene00010794 | DLD1_FOR<br>DLD1_REV | CCATCCACATCCAACCCTCTC<br>GGGGATTTCGGGATTTCGGG | 170 |
| *fard-1* | WBGene00022200 | FARD1_FOR<br>FARD1_REV | GTCAGGTCTTCTTCATGATCCCC<br>TCGTCGAACTTGACAGTCGC | 189 |
| *fat-6* | WBGene00001398 | FAT6_FOR<br>FAT6_REV | CAAGAGGAGAGCAAGAAGATCCC<br>TCACGGTTTGCCATTTTGCC | 133 |
| *ftn-2* | WBGene00001501 | FTN2_FOR<br>FTN2_REV | TGAAGTTGCACTCGACAGCC<br>TCGTTGATAGATTTGACCTGCTCG | 97 |
| *gdh-1* | WBGene00014095 | GDH1_FOR<br>GDH1_REV | ATTGAGAAGATCACCCGCCG<br>ACAAGCCGAAGCATCTCTGTC | 171 |

| | | | | |
|---|---|---|---|---|
| *gpx-7* | WBGene00019846 | GPX7_FOR<br>GPX7_REV | AGGGTGGAACTCTTTTCGATGC<br>TTGTCGACGGACCAAATCTCG | 96 |
| *idh-1* | WBGene00010317 | IDH1_FOR<br>IDH1_REV | GAATCCAATTGCCTCCATCTTCG<br>AATGGCAAGATCCTTGGTGAGG | 154 |
| *idhb-1* | WBGene00007993 | IDHB1_FOR<br>IDHB1_REV | AACTTGGCTGCTGGACTGG<br>AGTTGGGTTGGCGATAGAGC | 135 |
| *msra-1* | WBGene00018393 | MSRA1_FOR<br>MSRA1_REV | AAGGAGTTGTGGTGACACGAG<br>ATCGCAGATTGGTATTGCTTCTTTC | 196 |
| *pdha-1* | WBGene00011510 | PDHA1_FOR<br>PDHA1_REV | ACCCCAGCTCAAGAGATTCG<br>TGCCAGTCAATGTTGCATAGAA | 133 |
| *trx-2* | WBGene00007099 | TRX2_FOR<br>TRX2_REV | CTCTGGCGACTTCAAAAATGACAC<br>CCTTGTCTTCCGTTTACCTTTTCC | 199 |
| *T25B9.9* | WBGene00012015 | T25B9.9_FOR<br>T25B9.9_REV | TCCTCGGAGACATTGAACACG<br>CAACACGCCATGAATCCTGC | 111 |
