## Supplementary figures and images for "Redox homeostasis in ferroptosis and aging: a causal role for *fard-1* and *dhs-25* in *Caenorhabditis elegans*"

### Figure S1 800dpi.tif

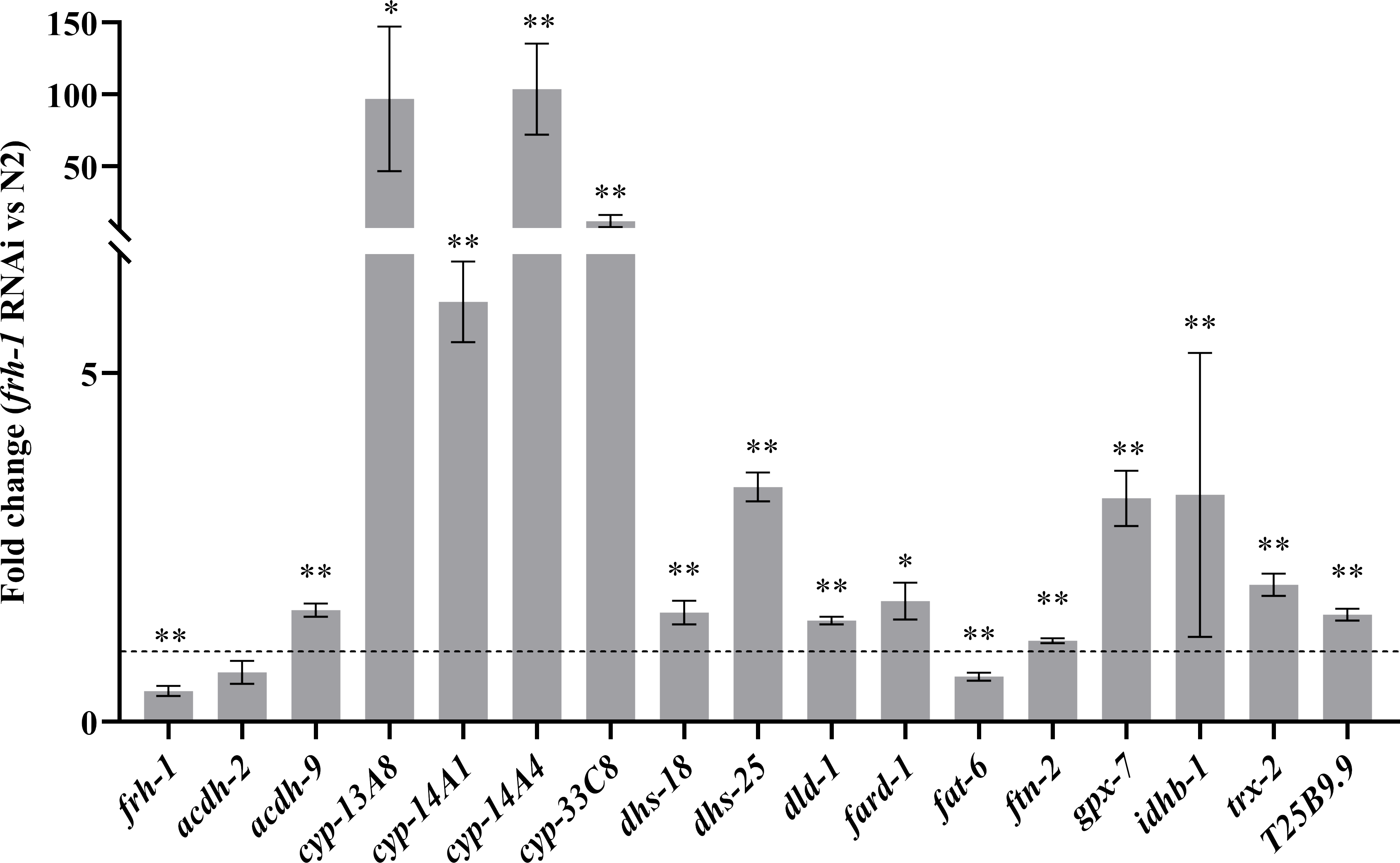

### Figure S3 800dpi.tif

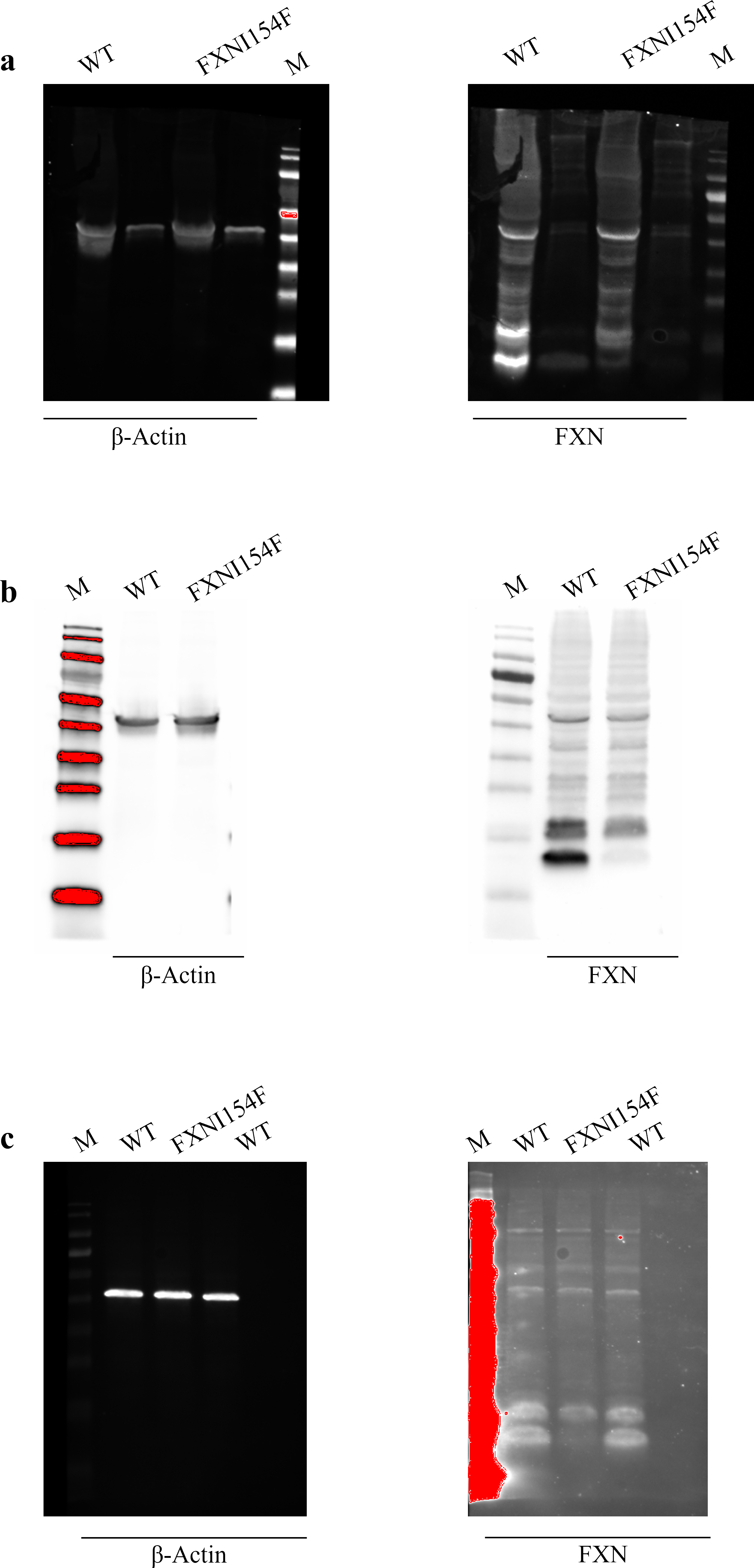
